# Environmental Factors Associated with Incidence and Distribution of *V. parahaemolyticus* and *V. vulnificus* in Chesapeake Bay, Maryland, USA: A three-year case study

**DOI:** 10.1101/2023.09.25.559351

**Authors:** Mayank Gangwar, Moiz Usmani, Yusuf Jamal, Kyle D. Brumfield, Anwar Huq, Avinash Unnikrishnan, Rita R. Colwell, Antarpreet S. Jutla

**Affiliations:** Geohealth and Hydrology Laboratory, Department of Environmental Engineering Sciences, University of Florida, Gainesville, FL, USA; Maryland Pathogen Research Institute, University of Maryland, College Park, MD, USA; University of Maryland Institute for Advanced Computer Studies, University of Maryland, College Park, MD, USA; Department of Civil, Construction, and Environmental Engineering, UAB School of Engineering, University of Alabama at Birmingham, AL, USA

**Keywords:** *Vibrio parahaemolyticus*, *Vibrio vulnificus*, *Vibrio*, environment, temperature, salinity, odds ratio, Chesapeake Bay

## Abstract

Members of the genus *Vibrio* are ecologically significant bacteria native to aquatic ecosystems globally, and a few can cause diseases in humans. Vibrio-related illnesses have increased in recent years, primarily attributed to changing environmental conditions. Therefore, understanding the role of environmental factors in the occurrence and growth of pathogenic strains is crucial for public health. Water, oyster, and sediment samples were collected between 2009 and 2012 from Chester River and Tangier Sound sites in Chesapeake Bay, Maryland, USA, to investigate the relationship between water temperature, salinity, and chlorophyll with the incidence and distribution of *Vibrio parahaemolyticus* (VP) and *Vibrio vulnificus* (VV). Odds ratio analysis was used to determine association between the likelihood of VP and VV presence and these environmental variables. Results suggested that water temperature threshold of 20°C or higher was associated with an increased risk, favoring the incidence of *Vibrio spp*. A significant difference in salinity was observed between the two sampling sites, with distinct ranges showing high odds ratio for *Vibrio* incidence, especially in water and sediment, emphasizing the impact of salinity on VP and VV incidence and distribution. Notably, salinity between 9-20 PPT consistently favored the *Vibrio* incidence across all samples. Relationship between chlorophyll concentrations and VP and VV incidence varied depending on sample type. However, chlorophyll range of 0-10 µg/L was identified as critical in oyster samples for both vibrios. Analysis of odds ratios for water samples demonstrated consistent outcomes across all environmental parameters, indicating water samples offer a more reliable indicator of *Vibrio spp.* incidence.

**Importance:** Understanding the role of environmental parameters in the occurrence of *Vibrio* species posing significant public health risks and economic burdens such as *Vibrio parahaemolyticus* and *Vibrio vulnificus* are of paramount importance. These aquatic bacteria are responsible for various human diseases, including gastroenteritis and wound infections, which can be severe and sometimes fatal. Recent observations suggest that certain environmental conditions may favor the growth of *Vibrio*, leading to more severe disease outcomes. By investigating the environmental factors that influence the occurrence of *Vibrio parahaemolyticus* and *Vibrio vulnificus*, the need to gain insights into the favorable ranges of environmental variables is apparent. The significance of this research is in identifying the favorable ranges of environmental and ecological factors, which holds the potential to provide an aid in the intervention and mitigation strategies through the development of predictive models, ultimately enhancing our ability to manage and control diseases caused by these pathogens.

## Introduction

*Vibrio spp.* are naturally occurring bacteria that play an ecologically significant role in the aquatic environment and have been shown to be associated with crustaceans and zooplankton. *Vibrio spp.* thrive in warm water with moderate salinity and their incidence is significantly influenced by environmental factors (1). Some *Vibrio* species are known to cause disease in humans(2), most notably *Vibrio cholerae,* the etiological agent of cholera and *Vibrio vulnificus* and *Vibrio parahaemolyticus*, which are associated with gastrointestinal illness and wound infections respectively (3). Recent studies have highlighted the global rise of *Vibrio* illnesses in humans, primarily attributed to changing environmental conditions and a poleward spread of pathogenic non-cholera *Vibrio spp*. (4, 5), which can pose significant public health risks (6), even in developed countries such as the United States (7–9). In the United States, *Vibrio* bacteria cause an estimated 80,000 illnesses and 100 deaths every year (9, 10). The diseases reported are often those associated with the consumption of raw or undercooked seafood (11–13). Since, *Vibrio spp.* are autochthonous to aquatic environments, especially coastal waters, and commonly concentrated in filter feeders, such as oysters and shellfish, which are often consumed raw, they offer an additional route of transmission to humans (14–16). Over the last several years, there has been an increase in the number of cases of *Vibrio*-related disease in United States (8, 9), often linked to the consumption of raw or undercooked seafood (11–13). This association most likely reflects that shellfish act as a reservoir for the bacteria, with *Vibrio spp.* detected in freshly harvested oyster meat and shellstock (11, 17, 18). Currently, in the United States, vibriosis causes an estimated 80,000 illnesses and 100 deaths every year (9, 10). In 1997 and 1998, there were four outbreaks involving more than 700 cases in the Gulf Coast, Pacific Northwest, and Atlantic Northeast regions, all of which were associated with the consumption of raw oysters.

Because oysters are filter feeders, they are able to concentrate bacteria, especially *Vibrio* spp., to such an extent that the concentration of bacteria in oysters will be much higher than that in the surrounding water (9) (8, 19, 20). The oyster, therefore, serves as a reservoir for *Vibrio spp.*, and also as a protective niche when environmental conditions become too harsh for growth and reproduction of the bacteria, e.g., winter temperatures, or as “passive concentrators”. Filter feeders such as oysters concentrate bacteria, including *Vibrio*, from the surrounding water (14, 21).

Chesapeake Bay, the largest estuary in the U.S., is located in the Mid-Atlantic region, where salinity and nutrient concentrations vary both seasonally and with the influx of freshwater (22). Notably, Chesapeake Bay is known for seafood production, especially blue crabs, clams, and oysters and is home to a multibillion-dollar commercial seafood industry providing seafood and oysters across the globe (23, 24). Hence, understanding the environmental influence on the distribution and growth of important *Vibrio spp.* is critical, considering the risk to human health and the economy. Previous work has shown that the incidence and distribution of *V. parahaemolyticus* and *V. vulnificus* is linked to environmental factors, including water temperature, salinity, and geographic location (16, 18, 25–27), dissolved oxygen (28–30), chlorophyll (31, 32), and plankton (33–35). A few of the earlier studies reported an association of chlorophyll, water temperature and salinity with the incidence of *V. parahaemolyticus* and *V. vulnificus*. The objective of this study was to determine the most favorable environmental parameters for likelihood of presence of these two vibrios in Chesapeake Bay and related bodies of water.

Several studies have examined the relationship between environmental parameters and the presence and abundance of *Vibrio* bacteria. Table 1 summarizes the known relationships of environmental parameters based on studies conducted in various geographical regions of the world. In the case of *V. vulnificus*, its abundance peaks when coastal water temperatures are greater than 19 °C (36) but often decreases with high temperature (37). During colder months of the year when coastal temperatures are less than 5°C, *V. vulnificus* is either not detected or detected only by employing molecular probes (20, 38, 39). Salinity has been found to be an important determinant of *V. vulnificus* abundance (10, 40), implying that vibrios respond to a favorable range of salinity, although a few exceptions were noted (Table 1) where a positive or no association was noted (40, 41).

**Table 1.**
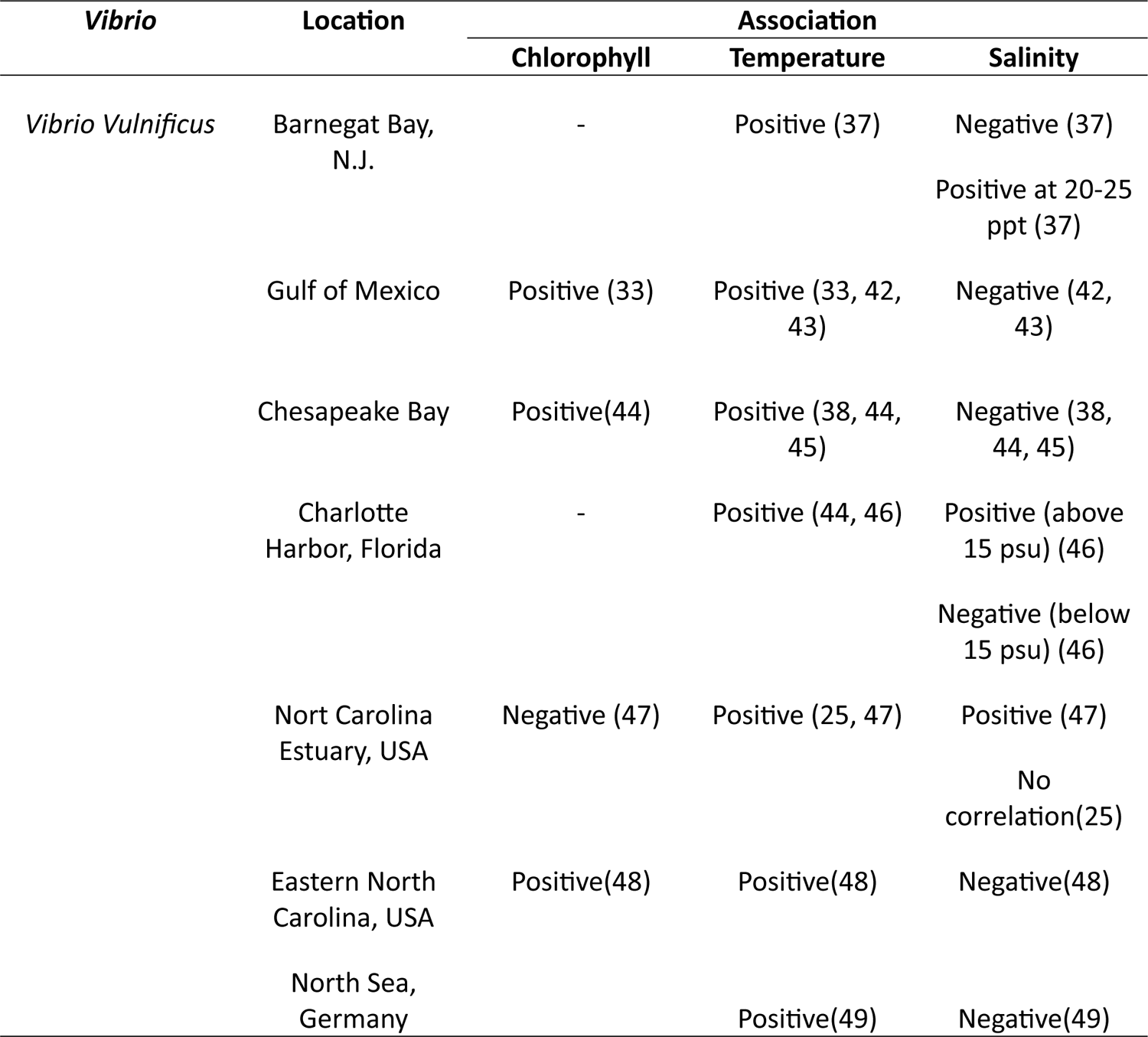
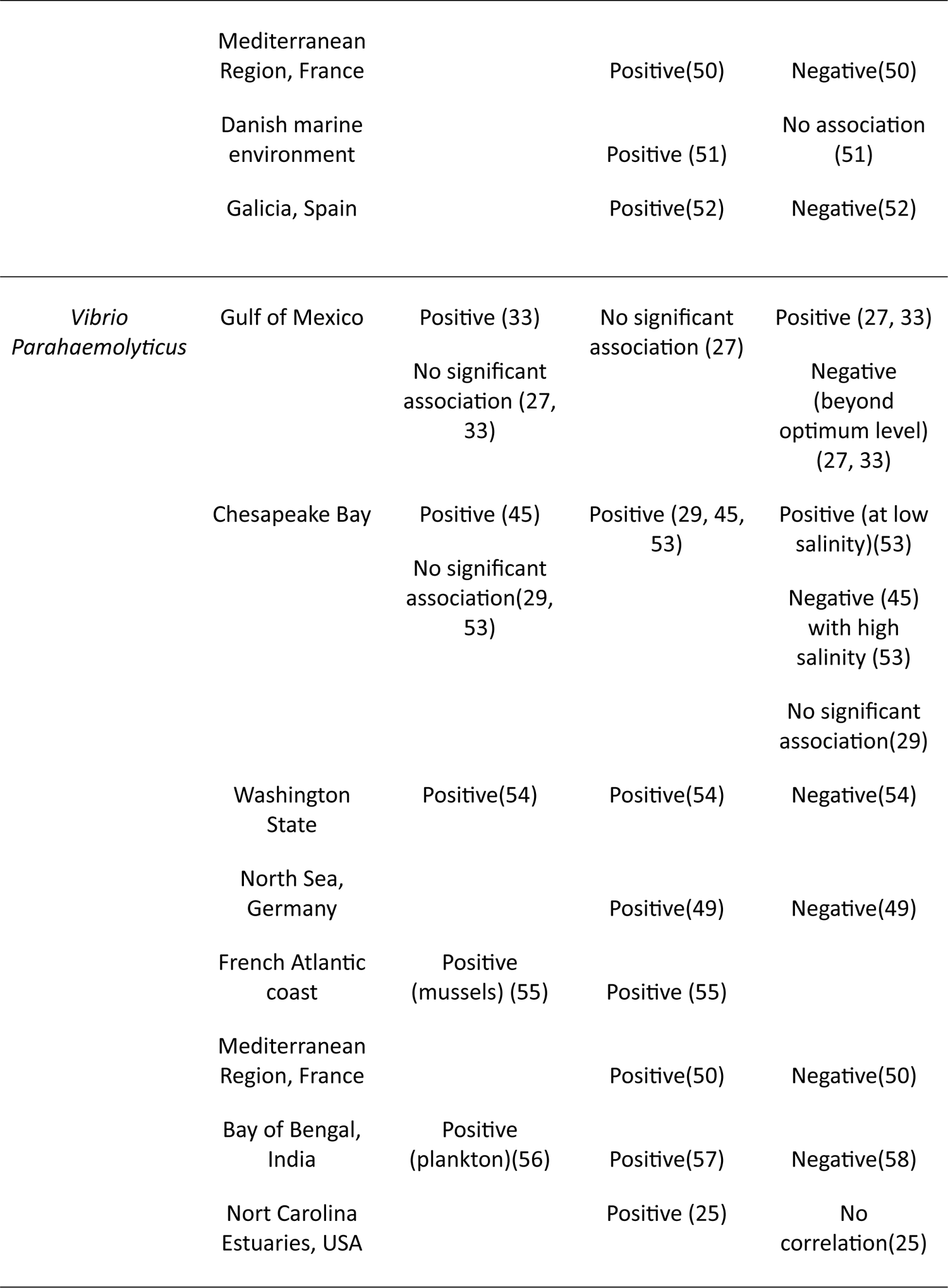
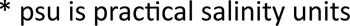
Environmental factors correlated with Vibrio spp. in different geographic locations.

| ***Vibrio*** | **Location** | **Association** |  |  |
| --- | --- | --- | --- | --- |
|  |  | **Chlorophyll** | **Temperature** | **Salinity** |
| *Vibrio Vulnificus* | Barnegat Bay, N.J. | - | Positive (37) | Negative (37)<br>Positive at 20-25 ppt (37) |
|  | Gulf of Mexico | Positive (33) | Positive (33, 42, 43) | Negative (42, 43) |
|  | Chesapeake Bay | Positive(44) | Positive (38, 44, 45) | Negative (38, 44, 45) |
|  | Charlotte Harbor, Florida | - | Positive (44, 46) | Positive (above 15 psu) (46)<br>Negative (below 15 psu) (46) |
|  | Nort Carolina Estuary, USA | Negative (47) | Positive (25, 47) | Positive (47)<br>No correlation(25) |
|  | Eastern North Carolina, USA | Positive(48) | Positive(48) | Negative(48) |
|  | North Sea, Germany |  | Positive(49) | Negative(49) |
|  | Mediterranean Region, France |  | Positive(50) | Negative(50) |
|  | Danish marine environment |  | Positive (51) | No association (51) |
|  | Galicia, Spain |  | Positive(52) | Negative(52) |
| *Vibrio Parahaemolyticus* | Gulf of Mexico | Positive (33)<br>No significant association (27, 33) | No significant association (27) | Positive (27, 33)<br>Negative (beyond optimum level) (27, 33) |
|  | Chesapeake Bay | Positive (45)<br>No significant association(29, 53) | Positive (29, 45, 53) | Positive (at low salinity)(53)<br>Negative (45) with high salinity (53)<br>No significant association(29) |
|  | Washington State | Positive(54) | Positive(54) | Negative(54) |
|  | North Sea, Germany |  | Positive(49) | Negative(49) |
|  | French Atlantic coast | Positive (mussels) (55) | Positive (55) |  |
|  | Mediterranean Region, France |  | Positive(50) | Negative(50) |
|  | Bay of Bengal, India | Positive (plankton)(56) | Positive(57) | Negative(58) |
|  | Nort Carolina Estuaries, USA |  | Positive (25) | No correlation(25) |

\* psu is practical salinity units

It is evident that some significant relationship exists for chlorophyll (phytoplankton), water temperature and salinity with *Vibrio* across various geographies. However, this also highlights a limitation in comprehensively utilizing existing data sources. This study aims to analyze the environmental parameters that influence the occurrence and abundance of *Vibrio spp.* in Chesapeake Bay using samples collected between 2009 and 2012. The study specifically focuses on determining the association of environmental parameters, such as chlorophyll, water temperature, and salinity, with the abundance of *V. parahaemolyticus* and *V. vulnificus*. However, the objective of this study is to identify the dominant threshold relationships of all three environmental variables and the incidence and occurrence of *V. parahaemolyticus* and *V. vulnificus*.

## Results

### Site evaluation

Each sampling location was first characterized, with respect to environmental parameters recorded at the time of each sampling event. Across all sampling events, pH, dissolved oxygen (DO), and water temperature ranges were relatively similar between the two sites (Fig 1). Fig 1 shows box plots summarizing the distribution of environmental parameters across sampling sites, with the center bar as the median of each group and the whiskers representing 1.5 times the interquartile range (IQR). However, there were notable differences in the salinity, conductivity, and total dissolved solids (TDS) profiles. At TS, salinity (min = 7.3, max = 19.1, median = 13.8 parts per thousand [PPT]), conductivity (min = 11.3, max = 29.5, median = 22.9 milli-siemens per centimeter [mS/cm]), and TDS (min = 10.8, max = 18.9, median = 15.1 grams per liter [g/L]) were significantly higher (P < 0.05) compared to CR (salinity: min = 3.2, max = 12.5, median = 8.4 PPT; conductivity: min = 5.8, max = 21.0, median = 14.61 mS/cm; TDS: min = 3.7, max = 13.4, median = 9.6 g/L). On the other hand, the Chl-a concentrations in the CR (min = 2.7, max = 41.39, median = 13.1 micrograms per liter [µg/L]) were slightly higher (P < 0.05) than those in the TS (min = 3.6, max = 32.7, median = 11.3 µg/L). In summary, the two sites exhibited similar pH, DO, and water temperature ranges, but differed significantly in salinity, conductivity, TDS, and Chl-a concentrations.

**Fig 1.**
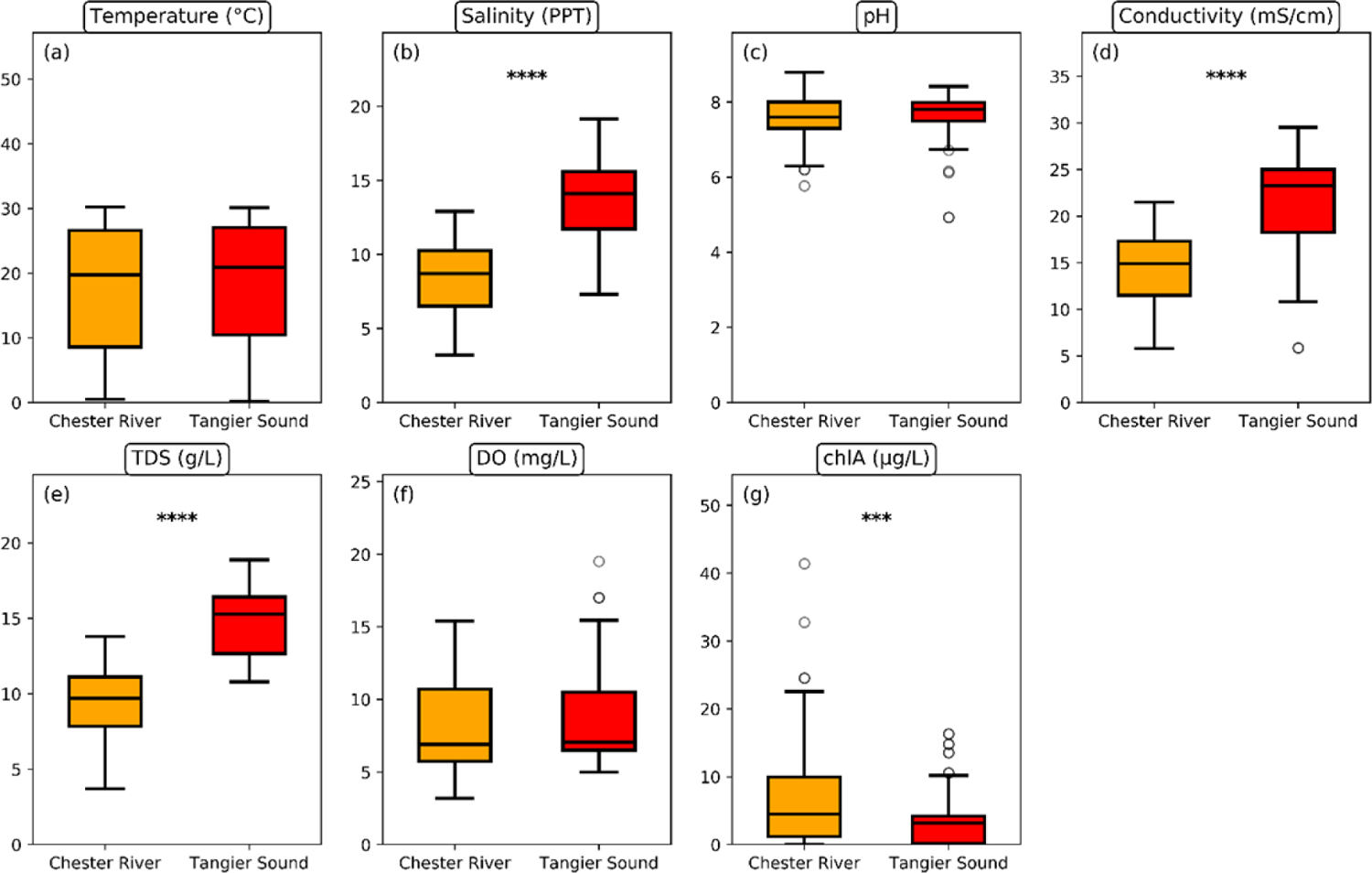
Distribution of environmental parameters (a) temperature in degree C; (b) salinity in parts per thousand (PPT); (c) pH; (d) conductivity in milli-Siemens per meter (mS/m); (e) total dissolved solids in grams per liter; (f) dissolved oxygen in milligrams per liter (mg/L); and (g) chlorophyll-a in micrograms per liter by sampling location. Any outlier values are represented by additional circles. The significant P values, determined using the t-test method, are displayed as follows: (**): ≤ 0.01, (***): ≤ 0.001, (****): ≤ 0.0001.

### Distribution and abundance of *V. vulnificus* and *V. parahaemolyticus*

*V. parahaemolyticus* and *V. vulnificus* incidence (presence/absence) and abundance in the different sample types were determined using DNA colony hybridization (DPCH). The maximum number of culturable *Vibrio* was as high as 11150 CFU/ml; minimum, maximum, and median numbers are indicated in Table 2.

**Table 2.** Summary of Environmental Parameters and Vibrio densities in different samples and sites.

| Sample | Site | No. of Samples | Range of SST (°C)<br>(median) | Range of Chl (µg/L)<br>(median) | Range of salinity (PPT)<br>(median) | Range of no. of CFU/ml or g for respective DPCH result (median) |  |
| --- | --- | --- | --- | --- | --- | --- | --- |
|  |  |  |  |  |  | tlh | vvh |
| Water | TS | 54 | 0.1 - 30.14<br>(21.48) | 0.02 - 10.19<br>(3.05) | 7.3 - 19.1<br>(13.77) | 0 - 52 (0.5) | 0 - 11150<br>(85) |
|  | CR | 57 | 0.5 - 30<br>(19.75) | 0.06 - 24.54<br>(3.2) | 3.9 - 12.9<br>(8.75) | 0 - 740<br>(1.25) | 0 - 80 (2.75) |
| Oyster | TS | 54 | 0.5 - 29.81<br>(20.875) | 0.04 - 16.31<br>(3.15) | 8.2 - 19.16<br>(14.14) | 0 - 1555<br>(11.5) | 0 - 145<br>(1.75) |
|  | CR | 57 | 0.5 - 30<br>(19.75) | 0.06 - 24.54<br>(3.2) | 3.9 - 12.9<br>(8.75) | 0 - 10300<br>(15) | 0 - 2430<br>(28.25) |
| Sediment | TS | 54 | 0.5 - 29.81<br>(20.875) | 0.04 - 16.31<br>(3.15) | 8.2 - 19.16<br>(14.14) | 0 - 2000<br>(15) | 0 - 1220 (16) |
|  | CR | 57 | 0.5 - 30<br>(19.75) | 0.06 - 24.54<br>(3.2) | 3.9 - 12.9<br>(8.75) | 0 - 10300<br>(15) | 0 - 2430<br>(28.25) |

DPCH detected both *V. vulnificus and V. parahaemolyticus* in all sample types. However, the detection rates and incidence of these bacteria showed significant variation, depending on factors such as temperature, salinity, and chlorophyll concentration (refer to supplementary material figures S1, S2 and S3). Additionally, there was an overall seasonal variation in detection rates. Among the samples, *V. parahaemolyticus* was detected most frequently in oysters (65.1%), followed by sediment (63.9%) and water (57.3%). Conversely, *V. vulnificus* was most frequently detected in sediment (75.7%), followed by water (72.1%) and oysters (69.4%). These findings highlight the importance of considering various environmental factors and sample types when studying the prevalence and distribution of *V. parahaemolyticus* and *V. vulnificus* in the Chesapeake Bay region.

### Seasonality of *Vibrio*

The MPN of the relevant genetic markers was compared with the time series of water temperature to visualize the effect of seasonality (Fig 2). This information provides both the timing and intensity of *V. parahaemolyticus* and *V. vulnificus*. In Chesapeake Bay, *Vibrio* species were detected at highest number in oyster samples during the warm summer to fall months (May to August) and in the fall (September to November). During the fall months the genes *vvh* and *tlh* were detected in significant numbers, with the highest levels observed in September and October. In May, *Vibrio parahaemolyticus*, reaching a peak of 52 CFU/gm, was detected in sediment samples using the *tlh* gene. The highest number of *V. vulnificus* (*vvh* gene) was observed in sediment samples in November, with a count of 177 CFU/gm. Water samples revealed that *Vibrio spp.* were primarily detected during the summer, with the *tlh* gene number peaking in June at 32 CFU/ml. The *vvh* gene was detected throughout the summer months, with the highest numbers observed in July at 80 CFU/ml.

**Fig 2.**
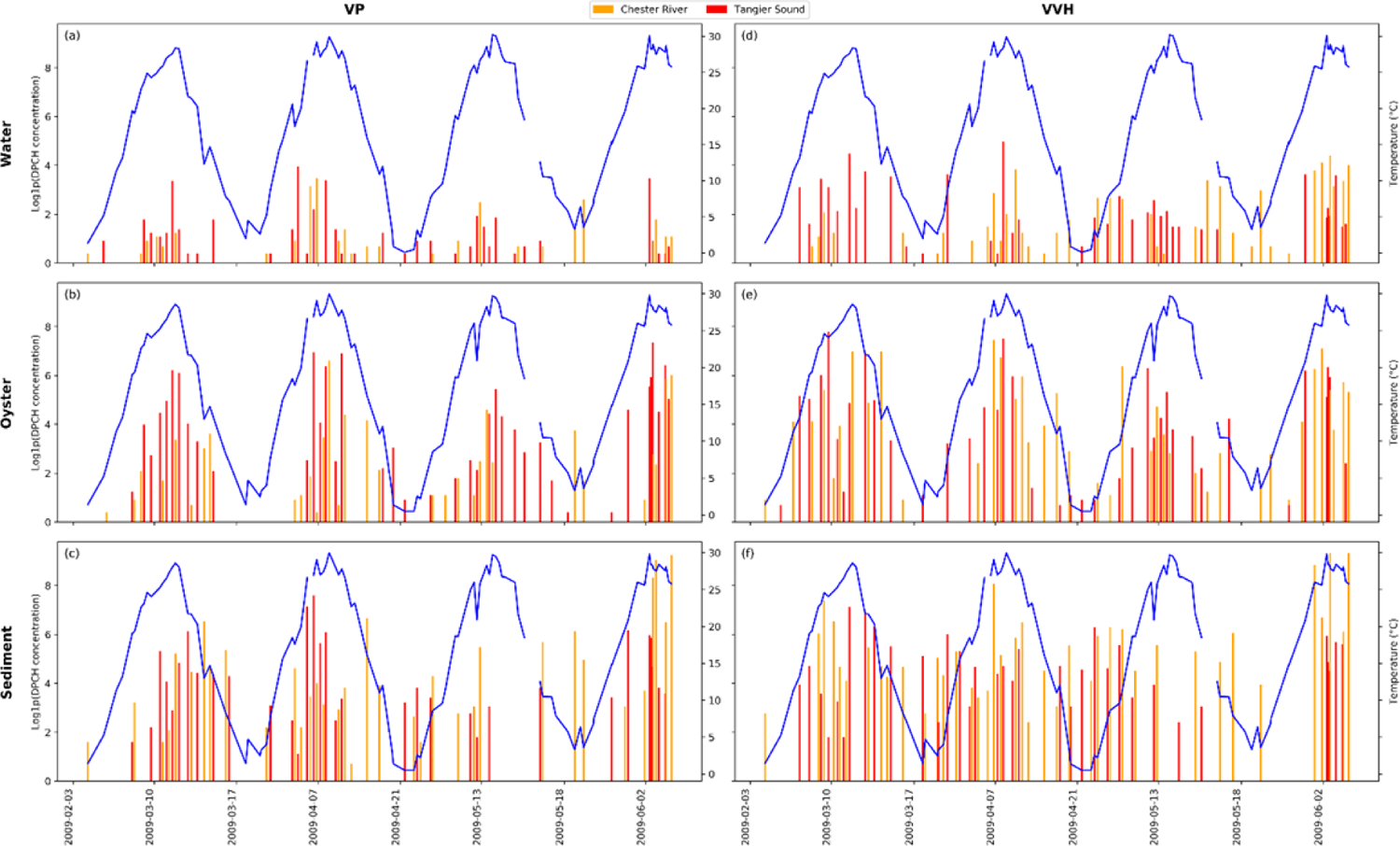
Bar plot showing population incidence of V. parahaemolyticus in (a) water column, (b) oyster, (c) sediment samples and V. vulnificus (d) water column, (e) oyster, (f) sediment samples, quantified by tlh and vvh markers employing DPCH. The left axis displays a bar plot of log1p(DPCH) results for total Vibrio parahaemolyticus (TLH) and Vibrio vulnificus (VVH), right axis shows water temperature recorded during sample collection.

### Analysis for identification of threshold environmental variables

The odds ratio analysis, a statistical metric used to evaluate the likelihood that an event (*Vibrio* incidence) will occur under a given condition to the likelihood that it will occur in the absence of a given condition, was employed to determine threshold values of environmental variables. The presence of *Vibrio spp.* was examined with respect to environmental parameters, namely water temperature, salinity, and chlorophyll concentration. Based on variations in environmental parameters, the odds ratio analysis enabled a quantitative assessment of the likelihood of the presence of *Vibrio spp.* The OR heatmaps are shown in Fig 3A and 3B for VP and VV respectively. The x-axis of the heatmap represents a relative unit of a variable. The y-axis represents the different environmental variables. The heatmap shows an OR greater than one, indicating a higher likelihood of VP presence in a sample compared to a reference. Colors vary based on odds ratio magnitude, with darker colors representing higher values.

**Fig 3.**
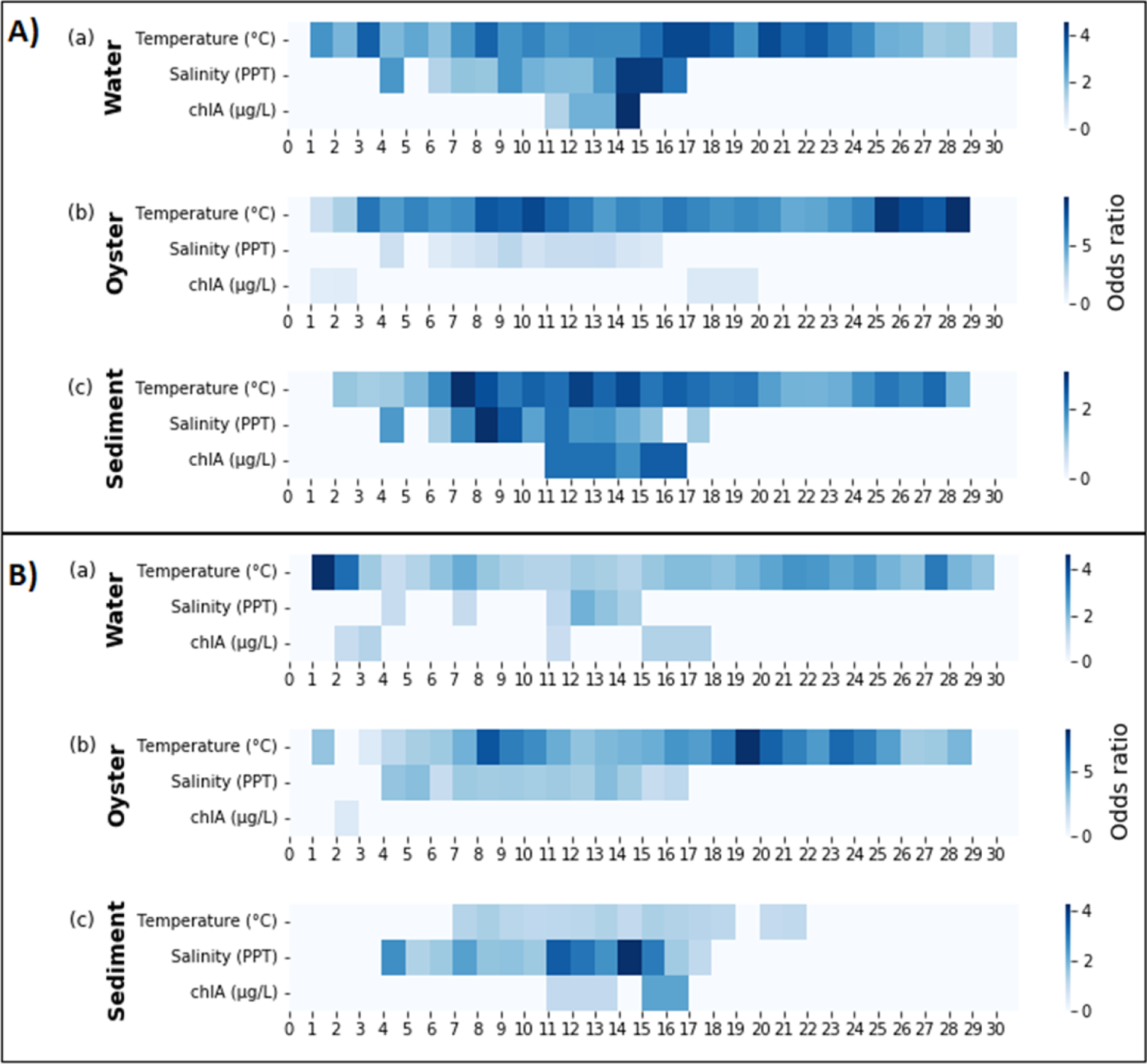
Heatmap of odds ratio for A) VP, and B) VV incidence; in (a) water, (b) oyster, and (c) sediment samples (x axis-relative unit of variable). Only odds ratios greater than 1 are shown.

The temperature results suggest that the incidence of VP and VV in the oyster samples is more probable across a wide temperature spectrum, even at lower temperatures (as shown in Fig 3A (b) and 3B (b)). This implies that both *Vibrio* species persist in oysters even in colder climates. In sediment samples, VP (Fig 3A. (c)) demonstrated higher odds than VV (Fig 3B (c)). The OR was higher for water samples at temperatures above 15°C. OR for extreme water temperatures was occasionally greater than one, a byproduct of the small number of samples collected at these temperatures, hence disregarded. The sediment samples did not yield significant odds for VV across any of the temperature ranges (Fig 3B. (c)).

For salinity, VP incidence in oysters (Fig 3A (b)) yielded a lower odds ratio compared to the water and sediment samples (Fig 3A (a) and 4A(c)). However, the salinity range corresponding to an odds ratio above one was consistent at 4 PPT to 17 PPT for all sample types. The VV odds ratio, with respect to the salinity of the water was low, compared to that of the oyster and sediment samples. In general, the odds ratio for salinity was higher for VP than VV.

The odds ratios for chlorophyll-a were high for water and sediment for VP (Fig 3A (a) and 3A(c). Chlorophyll did not reveal significance with respect to oysters for VV (Fig 3A (b)). For water and sediment, the latter showed a broader range of chlorophyll concentrations with a higher odds ratio for VP than water. For VV, OR values of chlorophyll for water and sediment were above one (Fig 3B (a) and 3B (c)), suggesting an association of chlorophyll with VV.

These results indicate a potential association of the environmental parameters and incidence of *Vibrio spp*. However, a conclusively favorable range for these parameters cannot be established from the available data. To determine thresholds for the environmental variables, ORs were computed for a range of 10 units for each variable. The 10-unit range ensured meaningful comparison and encompassed a sufficient range of *Vibrio* incidence to observe the influence of each variable on ORs. The decision to use a 10-unit range for each variable in determining the threshold for the environmental parameters was motivated by the standard deviation for temperature (the only continuous time series available for the dataset), which was ∼10°C. Furthermore, this standardized the analysis, rendering it comparable across the variables with understanding that there may be limitations in the variability of values. The subsequent ranges were limited to the integer immediately following the maximum value observed for the environmental variable. Corresponding OR heatmaps for VP and VV are shown in Fig 4A and 4B, respectively.

**Fig 4.**
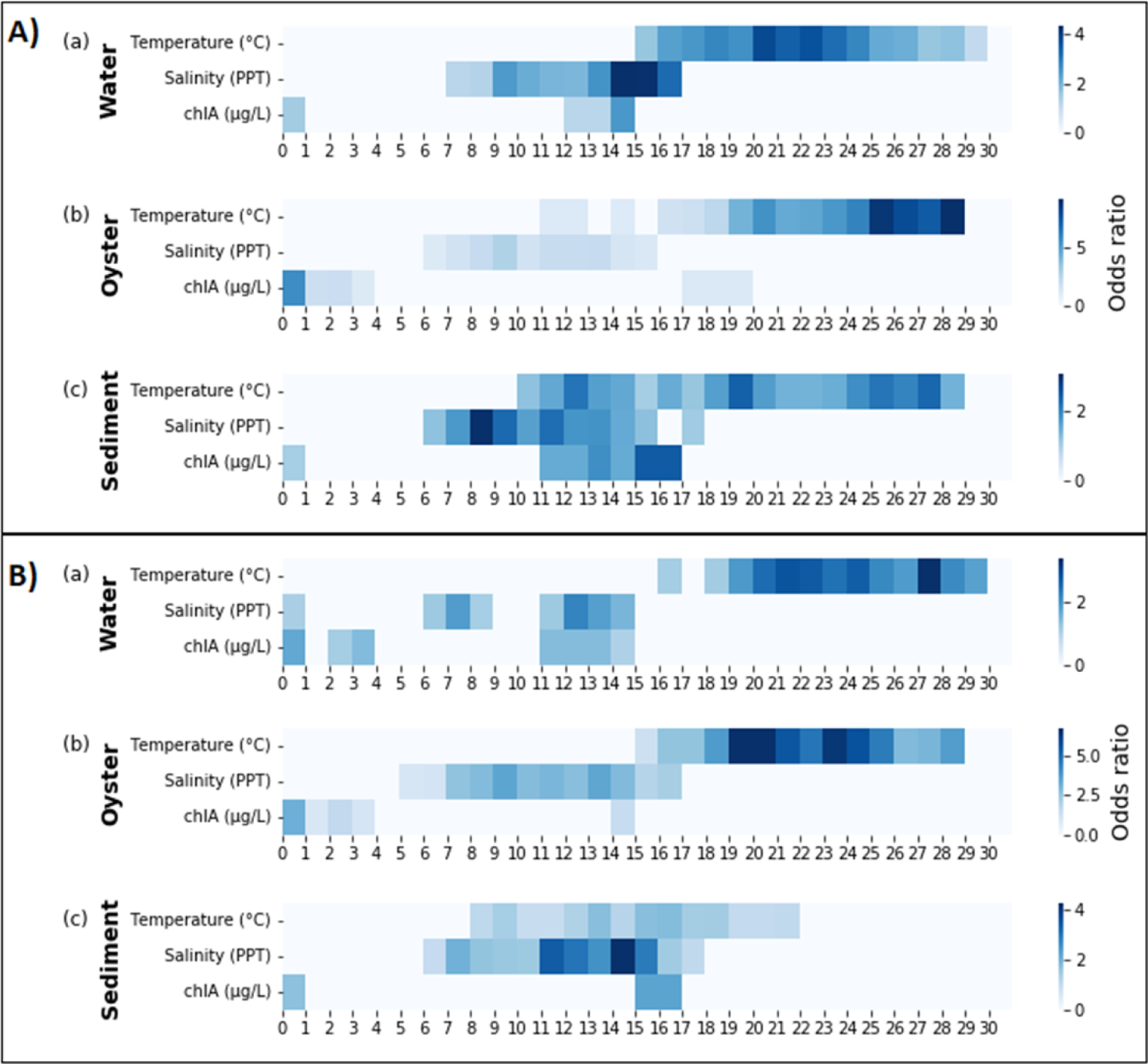
Heatmap of odds ratios (greater than one) in 10 units* of environmental variables for A) VP and B) VV incidence; in (a) water, (b) oyster, and (c) sediment samples. *The range variation is set at 10 units, meaning the scale of 0-1 corresponds to 0-10.

For water samples, temperatures exceeding 15°C were associated with higher occurrence odds for both VP and VV (Fig 4A (a) and 4B (a)). Similarly, oyster samples exhibited higher occurrence odds for both VP and VV when temperatures exceeded 16°C (Fig 4A (b) and 4B (b)). However, sediment samples exhibited a wider range of temperatures with higher odds for VP than VV (Fig 4A (c) and 4B (c)).

Salinity levels greater than 7 PPT were associated with higher *Vibrio spp.* occurrence across all samples. However, the odds were generally lower in oysters with respect to salinity. In the case of VV, two peaks were observed for OR in the water sample once after salinity of 7-17 PPT and second after 12-22 PPT. This pattern can be attributed to the difference in the salinity profile of the two sites, the Chester River and Tangier Sound. The more precise limits of determination remain inconclusive due to the small sample size.

It was observed that chlorophyll concentrations of 12-24 μg/L and 11-26 μg/L along with 0-10 μg/L were associated with increased odds of VP incidence in the water and sediment samples, respectively (Fig 4A). However, concentrations of 0-13 μg/L, as well as concentrations of 17-28 μg/L were found to be associated with chlorophyll concentration and VP incidence in the oyster samples (Fig 4A(c)). For VV, chlorophyll concentrations of 0-13 μg/L, as well as concentrations of 11-24 μg/L and 14-24 μg/L, were associated with increased odds of presence in water and oyster samples, respectively (Fig 4B a-b). Sediment samples showed higher odds at 0-10 and 15-26 μg/L, suggesting that in addition to low chlorophyll levels, higher chlorophyll concentrations, indicative of a higher phytoplankton biomass, may provide a more favorable environment for the *Vibrio*. This suggests that chlorophyll concentration influences the incidence of *Vibrio* in water and sediment differently than in oysters and may be due to the fact that oysters, as filter feeders, are able to concentrate *Vibrio* from the surrounding water regardless of the chlorophyll concentration. Alternatively, other factors, such as temperature or salinity, have a stronger influence on *Vibrio* incidence in oysters.

OR thresholds of greater than two and three were used to delineate favorable ranges or the threshold of environmental variables conducive to the incidence of *Vibrio* spp. An OR exceeding one implies a positive correlation between the exposure (environmental variable) and outcome (*Vibrio* occurrence). An OR surpassing 2 signifies a relatively stronger association, while an OR exceeding 3 denotes an even more robust association(59). By establishing a threshold of OR greater than 2 or 3, the aim is to pinpoint ranges of the environmental variables that significantly influence the incidence and distribution of *Vibrio*. These ranges are deemed favorable or threshold ranges as they demonstrate a markedly higher probability of *Vibrio* incidence relative to other ranges. These odds ratios were compared for predetermined thresholds (e.g., OR > 2 or 3). A range exceeding the threshold indicates a strong association between that specific range of the environmental variable and *Vibrio* incidence.

*V. parahaemolyticus* odds ratios greater than two were associated with specific ranges of environmental variables (Table 3(a)). For water samples, the temperatures between 16-31°C and the salinity of 9-20 PPT produce high OR. For oysters, the temperature range was 20-31°C and chlorophyll-a was 0-12 µg/L. For sediment, higher odds were obtained from as low as 12°C to 30.2°C max temperature, 8-20 PPT for salinity, and 15-26 µg/L for chlorophyll-a. When the odds ratio threshold increased to three, the temperature range for water narrowed to 20-31°C and salinity increased to 14-20 PPT (Table 3(b)). Hence the lower water temperature threshold of 20°C was found to have increased risk, with higher temperatures favor *Vibrio* incidence.

**Table 3.** Environmental variable range corresponding to a high odds ratio for Vibrio incidence.

|  | **Water** | **Oyster** | **Sediment** |
| --- | --- | --- | --- |
| **Temperature (°C)** | 16-31 | 18-31 | 12-22, 25-31 |
| **Salinity (PPT)** | 9-20 | 8-20 | 8-20 |
| **Chlorophyll (µg/L)** | 14-24 | 0-12 | 15-26 |

*(b) Odds ratio >3 for VP incidence*

|  | **Water** | **Oyster** | **Sediment** |
| --- | --- | --- | --- |
| **Temperature (°C)** | 20-31 | 19-31 | - |
| **Salinity (PPT)** | 14-20 | - | 8-18 |
| **Chlorophyll (µg/L)** | - | 0-10 | - |

*(c) Odds ratio >2 for VV incidence*

|  | **Water** | **Oyster** | **Sediment** |
| --- | --- | --- | --- |
| **Temperature (°C)** | 20-31 | 16-31 | - |
| **Salinity (PPT)** | 12-20 | 7-20 | 11-20 |
| **Chlorophyll (µg/L)** | - | 0-10 | 15-26 |

(d) Odds ratio >3 for VV incidence
|  | **Water** | **Oyster** | **Sediment** |
| --- | --- | --- | --- |
| **Temperature (°C)** | 27-31 | 18-31 | - |
| **Salinity (PPT)** | - | 9-20 | 11-20 |
| **Chlorophyll (µg/L)** | - | 0-10 | - |

*V. vulnificus* odds ratios above two corresponded to temperature ranges of 20-31°C in water and 16-30°C in oyster samples (Table 3(c)). The salinity ranges were 12-20 PPT in water, 7-20 PPT in oysters, and 11-25 PPT in sediments. Chlorophyll-a ranges were 0-10 µg/L for oysters and 15-26 µg/L for sediments. With an odds ratio threshold of three (Table 3(d)), the temperature range increased to 27-37°C for water and remained at 18-31°C for oysters. The salinity range remained 11-20 PPT for sediment and 9-20 PPT for oysters. These results highlight the significance of specific environmental parameters, notably temperature and salinity, influencing the incidence and distribution of *V. vulnificus* in the different sample types.

Odds ratios for water samples demonstrated uniform outcomes across all examined environmental parameters (temperature, salinity, and chlorophyll) which implies that water samples may serve as a more dependable and holistic indicator of *Vibrio* presence. This could be because these samples directly reflect the environment where *Vibrio* species inhabit and proliferate. Therefore, changes in environmental and ecological parameters such as temperature, salinity, and chlorophyll concentrations are likely to have a more immediate and direct impact on *Vibrio* presence in water samples compared to oyster and sediment samples.

The findings suggest that the incidence of *V. parahaemolyticus* and *V. vulnificus* in oyster samples is more probable across a wide temperature spectrum, even at lower temperatures. This could imply that both types of vibrios might be capable of surviving, and potentially thriving, in oysters even in colder climates. *V. vulnificus* exhibited a preference for the water column, while *V. parahaemolyticus* tended to favor the sediment. The higher abundance of *V. vulnificus* in the water column likely contributes to its elevated levels in oysters compared to *V. parahaemolyticus*.

## Discussion

Both the incidence and distribution of *V. parahaemolyticus* and *V. vulnificus* in the Chester River and Tangier Sound are high during the summer and fall seasons. The abundance of these bacteria was quantified using a molecular approach employing specific genetic markers (*tlh* and *vvh*) for the species. *V. vulnificus* was detected at high concentrations during the two seasons, June and October-November, and *V. parahaemolyticus* was detected during June-July and October-November. However, *V. vulnificus* was detected in higher numbers than *V. parahaemolyticus* in all of the samples at both sites.

Environmental factors such as chlorophyll and water temperature have been demonstrated to significantly influence the growth and distribution of *Vibrio* bacteria (47). The temperature impacts the growth of bacteria, and chlorophyll indicates the presence of nutrients that support the growth of phytoplankton, which can subsequently serve as a food supply for bacteria like vibrios(60). Salinity has previously been shown to play a crucial role in bacterial survival and virulence and provides vital information that can significantly affect the growth and spread of vibrios(35, 37, 53). It can alter the bacteria’s osmotic balance and metabolic processes, thereby influencing their survival and proliferation (50). The odds ratio results highlight the significant influence of specific environmental parameters on the presence of *V. parahaemolyticus* and *V. vulnificus* across different coastal ecosystems, namely the water column, oysters, and sediment. The findings suggest that there may be an optimal range of environmental and ecological variables: surface water temperature, salinity, and chlorophyll concentration favoring *Vibrio* bacteria.

The results provide additional evidence that water temperature is a major determinant of the occurrence and distribution of *Vibrio spp.* in the environment (16, 61, 62). It has been identified that surface water temperature is critical in influencing the growth and reproduction of *Vibrio spp.* The elevated temperatures are associated with an increased likelihood of *Vibrio* incidence, indicating that warmer months of the year are more conducive to the proliferation of *Vibrio spp.* It is found that the higher temperature range for the oyster samples, compared to water and sediment, is closely linked to heightened odds of detecting both *V. parahaemolyticus* and *V. vulnificus*. This reaffirms the notion that oysters serve as reservoirs for *Vibrio spp.* and their ability to harbor *Vibrio spp.* during colder temperature months. Interestingly, other environmental factors, namely salinity and chlorophyll, exert a greater influence on the incidence of *Vibrio spp.* in oyster samples, compared to water and sediment.

Salinity plays a significant role in the incidence and distribution of *Vibrio* species (37, 53). Distinct ranges of salinity levels were found to be associated with high odds ratios, suggesting that specific salinity conditions favor the growth and proliferation of VP and VV in water and sediment. Chlorophyll concentration, especially in oyster samples, also contributed to a favorable environment for these bacteria. Water samples showed consistent odds ratios for *Vibrio* incidence across all environmental parameters (temperature, salinity, and chlorophyll), suggesting that water is a more reliable indicator of the environmental impact on *Vibrio* incidence than oysters. In addition to physical and chemical variables associated with the incidence and distribution of *Vibrio spp.,* there are other factors which influence the incidence and distribution of VV and VP, such as hydrostatic pressure, since with greater depth of the water column a statistically significant increase in the number of *V. vulnificus* has been reported (47, 63). This paves the path for further investigations to better understand the relationship between *Vibrio spp.* and their host organisms, as well as the role of different environmental factors in this interaction.

Gaining a comprehensive understanding of the ecology of *Vibrio spp.* and various environmental factors that influence their occurrence and transmission is of great significance. This study underscores the significant influence of distinct environmental parameters on the incidence of *V. parahaemolyticus* and *V. vulnificus* in the Chesapeake Bay, findings that are extrapolatable to other coastal ecosystems, with reference to water, oysters, and sediment. Both VP and VV exhibited seasonal variation, with higher numbers observed during the warm summer to fall months. The incidence of VP and VV in oyster samples was found to be likely across a wide temperature spectrum, even at relatively low temperatures, with the implication that both VV and VP can endure in oysters during the cold months of the year. *V. vulnificus* was found to have significantly higher odds of occurrence in the water column, while *V. parahaemolyticus* tends to favor the sediment. The higher abundance of *V. vulnificus* in water is likely associated with its higher incidence in oysters, which are filter feeders, compared to V. parahaemolyticus.

The findings provide a platform for future research on the environmental factors that influence *Vibrio* bacteria. Other factors to be considered include anthropogenic activities. Climate variability, agriculture, and urban runoff, all of which can alter environmental conditions, especially nutrient input, that can affect the incidence and distribution of *Vibrio spp.* Such factors contributing to *Vibrio* dynamics and their implication for public health will need to be studied to understand the impact of convective processes on *Vibrio* dynamics.

## Material and Methods

### Site Description and Sample Collection

Methods for sample collection and processing have been described in detail previously (10). A summary relevant to the study reported here is as follows. Briefly, samples were collected at two stations, one in the Chester River (CR) and the other in Tangier Sound (TS) (Fig 5). The CR is a significant tributary, located in the northern part of Chesapeake Bay (39°05.09’N, 76°09.50’W) and is characterized by brackish water. In contrast, the TS is an inlet of the Atlantic Ocean farther south (38°10.97’N, 75°57.90’W). From June 2009 to August 2012, water, oyster, and sediment samples were collected at two locations in the CR and TS. A total of 110 samples, including water and oysters, were collected at these two stations, CR (n = 56) and TS (n = 54). During the warmer months of the year (June through August), sampling was performed twice each month and once each month from September through May. Water samples were collected from the surface (1 ft) and bottom (1 ft from the bottom) at each site.

**Fig 5.**
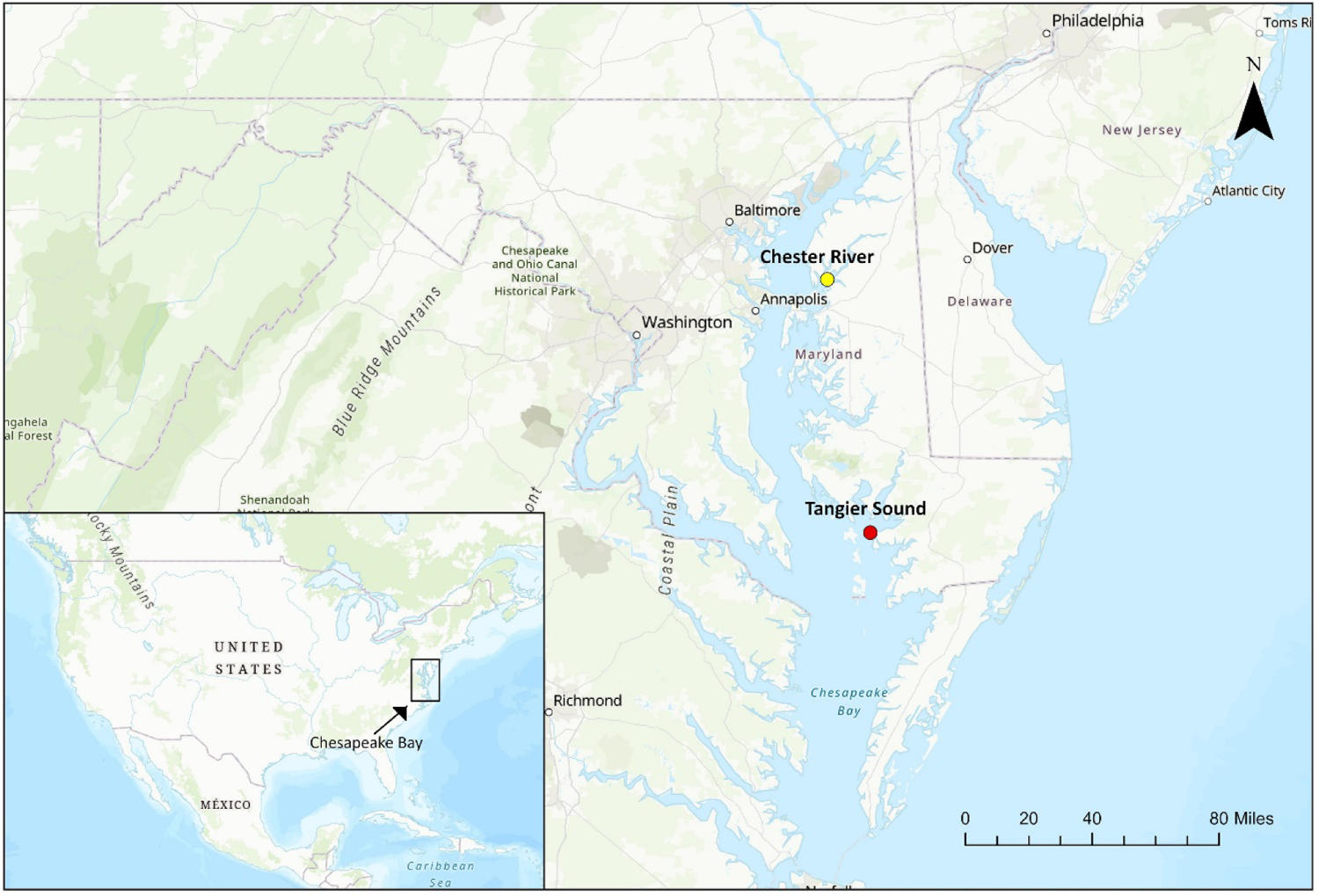
Chesapeake Bay sampling sites. The map illustrates the locations where samples were collected, the Chester River (yellow circle) and Tangier Sound (red circle). An inset map of the USA is provided to indicate the location of Chesapeake Bay. The scale bar on the map corresponds to the distance depicted.

### Environmental data

Water parameters, including water temperature, conductivity, salinity, TDS, chlorophyll-a, pH, dissolved oxygen, and turbidity were determined on site for samples collected at the surface (1 ft) and bottom (1 ft from the bottom) at each site, and conductivity measurements were made using a digital handheld conductivity meter (model 30-25FT; Yellow Springs Instruments, Yellow Springs, OH). At each site, 12 liters of water, 20-25 oysters, and 80-100 g of sediment (collected from oyster boxes or beds) were collected. Samples were kept on ice during transport to the laboratory at the University of Maryland, College Park and, upon arrival the samples were stored overnight at 15°C until processing the following morning.

Turbidity was measured at the site using a Secchi disk. TDS was measured in the laboratory using oven-drying while chlorophyll-a and other pigments were measured in methanol extracts using a Cary model 50 UV–visible-light spectrophotometer. Dissolved organic carbon analysis was done using a Shimadzu TOC-V CSN carbon analyzer equipped with an ASI-V autosampler (Shimadzu Scientific Instruments, Columbia, MD).

### Direct Plating and DNA Colony Hybridization

Quantification of *V*. *parahaemolyticus* and *V*. *vulnificus* populations was performed using the direct plating and DNA colony hybridization (DPCH) method targeting the thermolabile hemolysin (*tlh*) and *V. vulnificus* hemolysin, *vvhA* genes, respectively, as previously described (10).

### Statistical analysis

A descriptive analysis was performed to determine the incidence and distribution of vibrios with respect to environmental parameters. Variable input included the average number of colony forming units (CFU) observed across duplicate hybridizations, using water (CFU/ml), oyster (CFU/0.1 g), and sediment (CFU/0.1 g), for *V. parahaemolyticus* and *V. vulnificus*, along with temperature in degree C, salinity in part per thousand (PPT), chlorophyll-a in microgram per liter (µg/L). In the analysis, various tools were utilized, and their respective versions were recorded for accuracy and reproducibility. The tools employed include Pandas (version 0.24.2), NumPy (version 1.16.2), Matplotlib (version 3.0.3), Seaborn (version 0.11.2) and Scikit-learn (version 0.24.2). Additionally, Python version 3.6.8 was used as the programming language throughout the analysis. Proper references were consulted for each tool to ensure the validity of the results and to adhere to best practices in data analysis. The association between study variables was explored using univariate analysis. Bar plots were used to determine the distribution of the *Vibrio* genes (*tlh and vvh*). The scatter plots display the abundance of *Vibrio parahaemolyticus* and *Vibrio vulnificus* across each sample type (Fig S1, S2 and S3). Water, sediment, and oyster samples were used to explore the relative incidence of each of the *Vibrio* genes with respect to sample type.

Since the data are in the pulse format (0 and 1), Odds Ratios (OR) were used to determine associations (64, 65) among presence of vibrios and the corresponding environmental condition, indicating how closely an incident was linked to exposure by describing the ratio of presence odds given a variable threshold (65). The OR has been widely used in environmental studies to identify the thresholds at which the presence of a particular organism, such as *Vibrio*, is significantly related to environmental factors. OR compares probabilities of the likelihood that an event will occur in a given condition with respect to the likelihood that the event will occur in the absence of the given condition. Therefore, OR determines how likely an exposure will cause a particular occurrence, the likelihood of which will increase proportionally with the OR value, and OR values less than one indicate that the event has a lesser chance of happening given the exposure. To evaluate the association between events and pre-harvest environmental conditions of *Vibrio* (66, 67), we use the following equation:

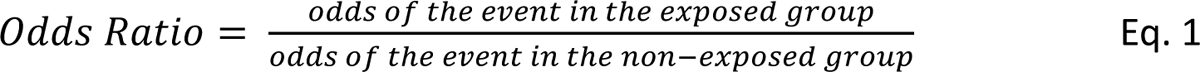

The OR value can be used to estimate the probability of *Vibrio* presence in a given specific environmental condition. The likelihood of *Vibrio* occurrence can be estimated by analyzing the association between *Vibrio* presence and environmental factors such as temperature, which can assist in evaluating the likelihood of *Vibrio* occurrence given specific environmental conditions. Since *Vibrio* detection between the two stations included in this study lacks substantiative differences, therefore data were pooled prior to OR analysis.

We conducted an analysis to examine in detail how specific ranges of the environmental variables were associated with the incidence of vibrios. Procedures to determine the range of an environmental variable that promotes *Vibrio* presence involved several steps. The data were divided into two categories: the first included observations where environmental variables were within the observed range (referred to as the “exposure” group), while the second included observations where the environmental variables were outside this range (“non-exposure” group). Within each category, the number of “successes” and “failures” was determined. A “success” was defined as the presence of *Vibrio*, while a “failure” indicated its absence. The odds of success in each group were then calculated as the ratio of the number of successes to the number of failures. Initially, the odds ratio was calculated for each value of the environmental parameter as the ratio of the odds in the exposure group to the odds in the non-exposure group. An OR greater than one suggested that the presence of *Vibrio* was more likely above the considered value for that environmental parameter, while an OR less than one suggested that it was less likely. By calculating the odds ratios for each value of the environmental variable, we were able to illustrate how the likelihood of *Vibrio* occurrence changes with varying values of the environmental variable. This reveals a compelling association with elevated odds ratios, indicating that samples falling within these specific ranges exhibit a substantially heightened incidence of *Vibrio*. This facilitated an in-depth exploration of the relationships between the environmental variable and *Vibrio* incidence. This was repeated for each environmental variable, allowing comparison of ORs to determine limits of threshold values, as well as environmental parameters having the greatest influence on the incidence of *Vibrio spp*. in the water, sediment and oyster samples. Finally, the range of environmental factors with consistently high odds ratios, indicative of a high risk for *Vibrio* incidence, was identified.

## Data availability

The data used in the current study have been deposited in a private GitHub repository and can be made available to the reviewer upon request. Interested reviewers can contact the corresponding author at **Error! Hyperlink reference not valid.** to access the data. Once the paper is accepted, the data will be made publicly accessible at https://github.com/GangwarMayank/Chesapeake-Bay-Vibrio-parahaemolyticus-and-Vibrio-vulnificus-occurrence-data.git.

## Authors’ contributions

**Conceptualization:** MG, MU, ASJ **; Data curation:** MG and KDB; **Formal Analysis:** MG, MU, YJ, KDB; **Funding acquisition:** ASJ, AH and RRC; **Methodology:** all authors; **Project administration:** ASJ, AH, and RRC; **Resources:** ASJ, AH, and RRC; **Software:** MG; **Supervision:** ASJ, AU, AH, and RRC; **Validation:** all authors; **Visualization:** MG, MU, YJ, and ASJ; **Writing-original draft:** MG, MU, YJ and KDB; **Writing-review and editing:** all authors; All authors read and approved the final manuscript.

## Acknowledgements

This research was supported by the National Institute of Environmental Health Sciences, National Institutes of Health (NIH) under award number R01ES030317A, and the National Science Foundation under award number OCE1839171 and CBET 1751854. The authors declare that they have no conflict of interest.

## Supplementary material

**Figure S1.**
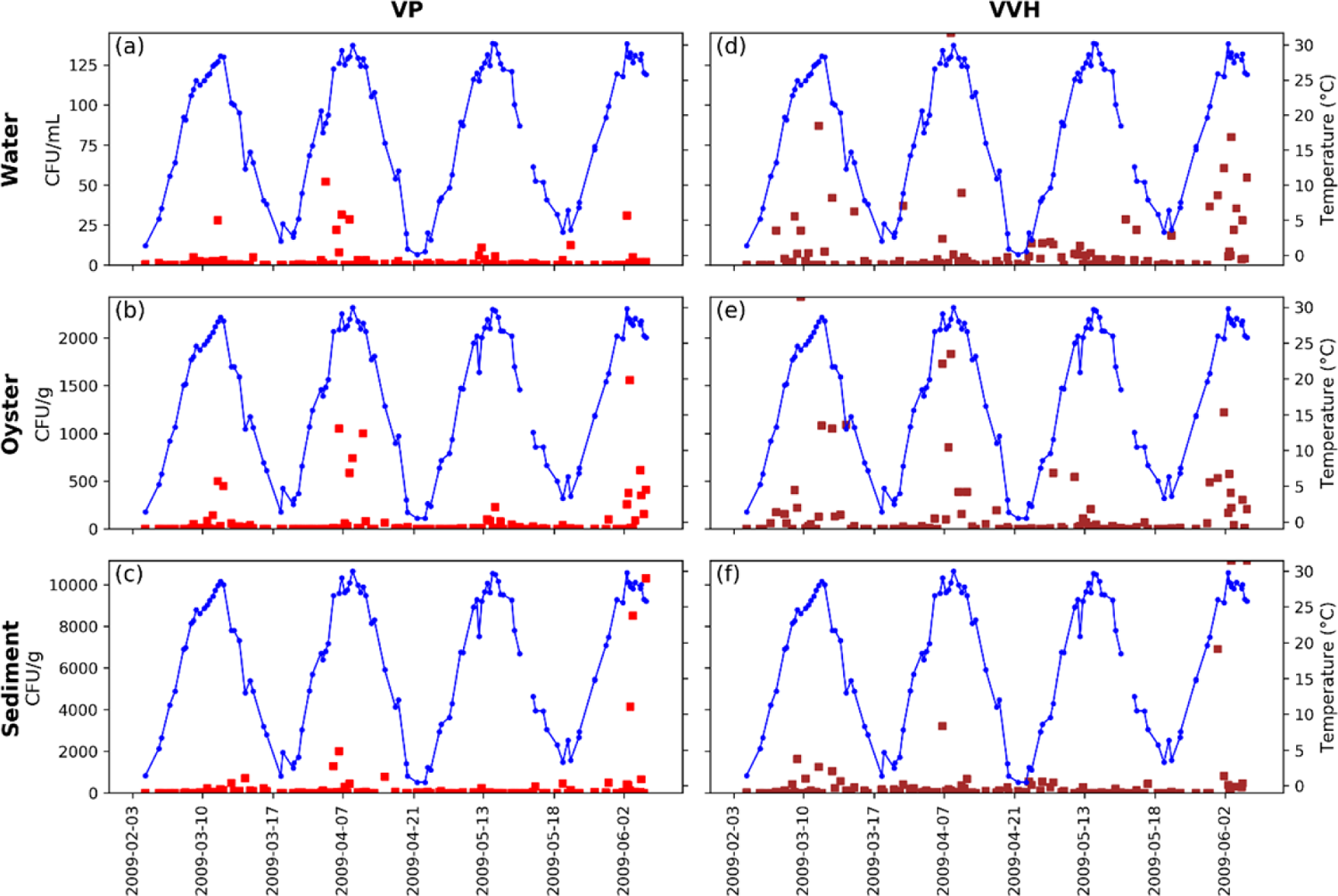
Incidence of V. parahaemolyticus in (a) water column, (b) oyster, (c) sediment samples and V. vulnificus (d) water column, (e) oyster, (f) sediment samples, quantified by tlh and vvh markers through DPCH vs temperature (°C). Red squares, density of tlh or vvh in number of CFU/ml (water) or CFU/g (oysters and sediment); blue circles, sea surface temperature in °C.

**Figure S2.**
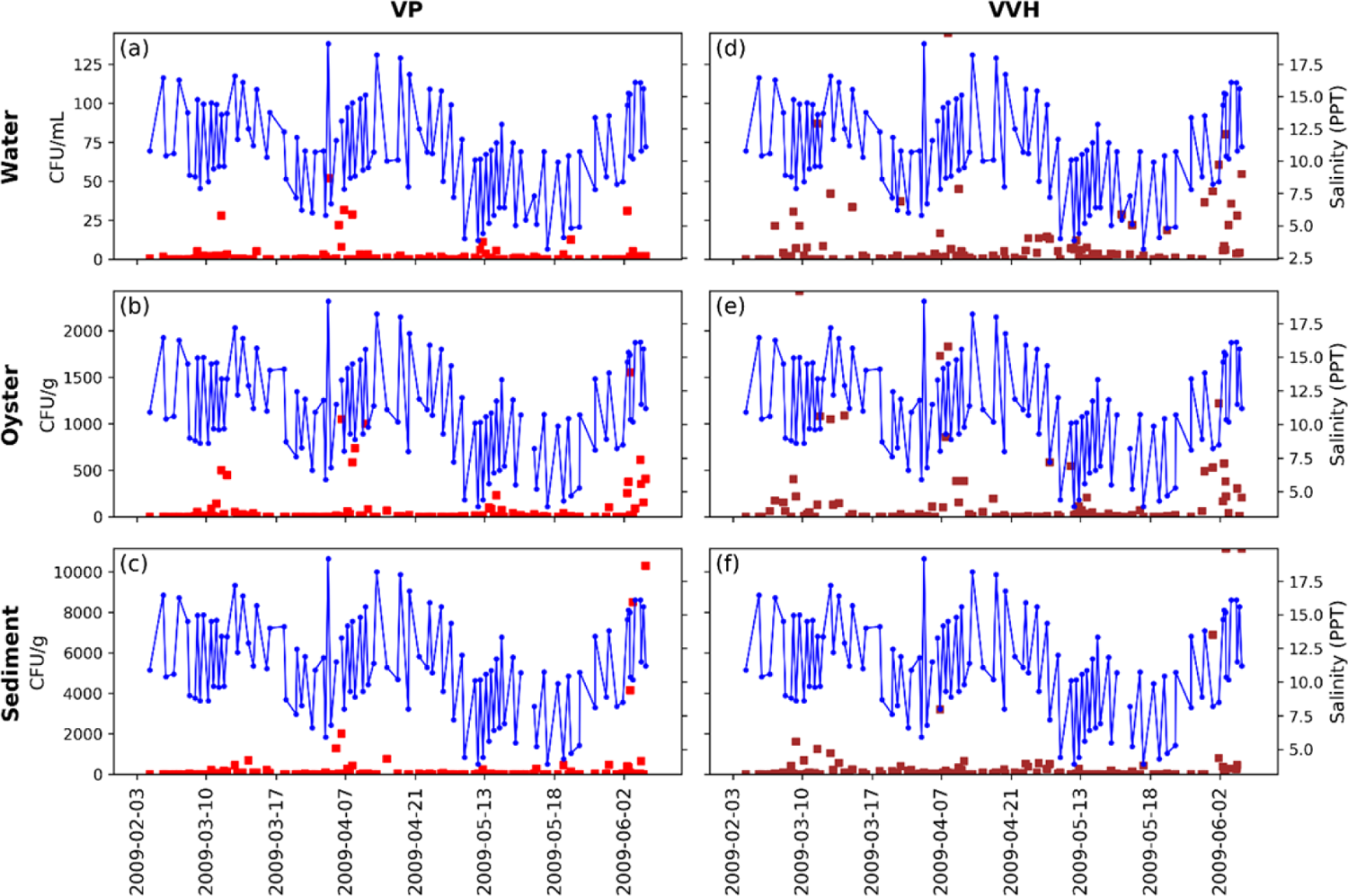
Incidence of V. parahaemolyticus in (a) water column, (b) oyster, (c) sediment samples and V. vulnificus (d) water column, (e) oyster, (f) sediment samples, quantified by tlh and vvh markers through DPCH vs salinity (PPT). Red squares, density of tlh or vvh in number of CFU/ml (water) or CFU/g (oysters and sediment); blue circles, salinity in PPT.

**Figure S3.**
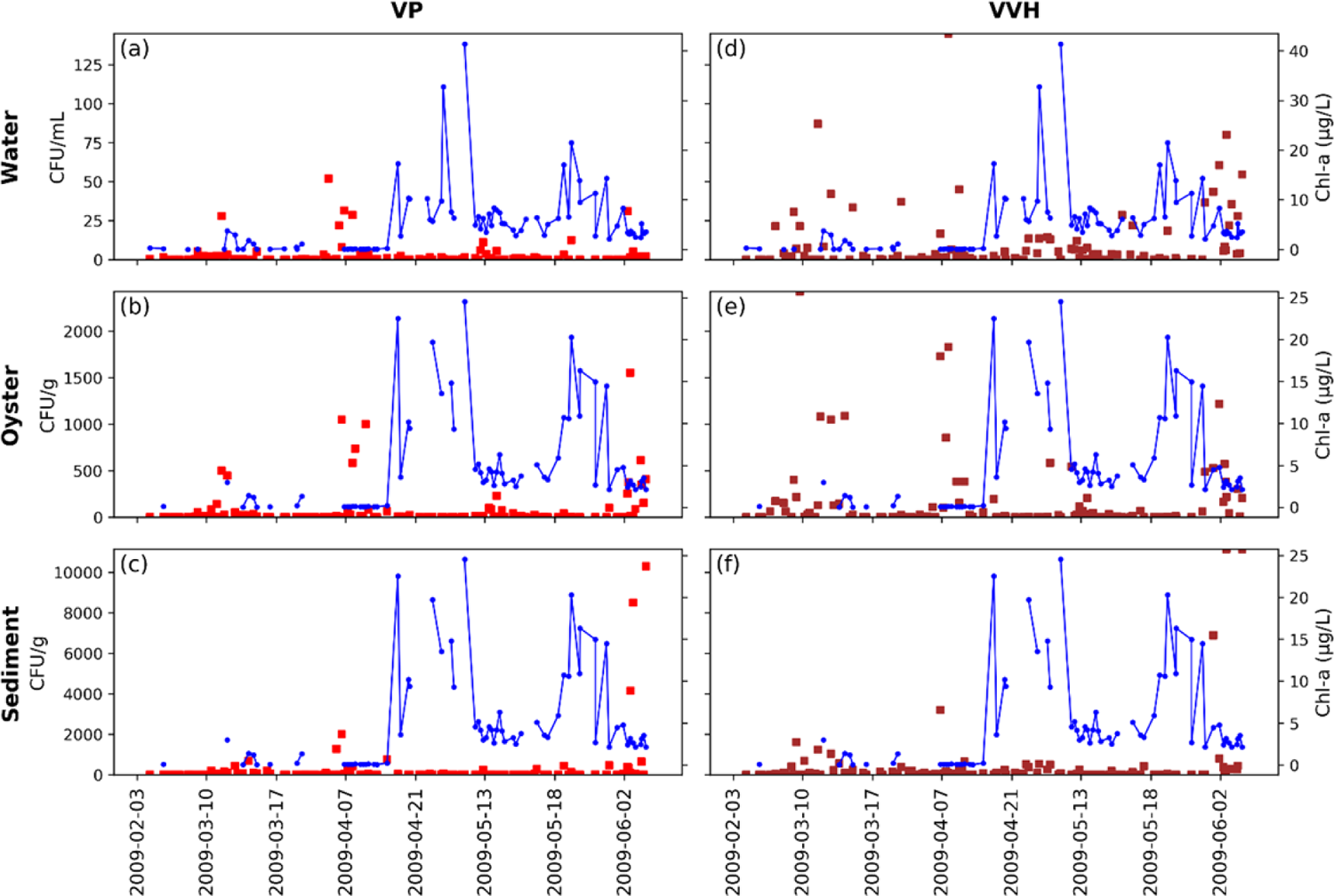
Incidence of V. parahaemolyticus in (a) water column, (b) oyster, (c) sediment samples and V. vulnificus (d) water column, (e) oyster, (f) sediment samples, quantified by tlh and vvh markers through DPCH vs chlorophyll concentration in µg/L. Red squares, density of tlh or vvh in number of CFU/ml (water) or CFU/g (oysters and sediment); blue circles, chlorophyll concentration in µg/L.

## References

1. Brumfield KD, Usmani M, Chen KM, Gangwar M, Jutla AS, Huq A, Colwell RR. 2021. Environmental parameters associated with incidence and transmission of pathogenic *Vibrio spp*. Environmental Microbiology 23:7314–7340.

2. Baker-Austin C, Trinanes J, Gonzalez-Escalona N, Martinez-Urtaza J. 2017. Non-Cholera Vibrios: The Microbial Barometer of Climate Change. Trends Microbiol 25:76–84.

3. Heng S-P, Letchumanan V, Deng C-Y, Ab Mutalib N-S, Khan TM, Chuah L-H, Chan K-G, Goh B-H, Pusparajah P, Lee L-H. 2017. Vibrio vulnificus: An Environmental and Clinical Burden. Front Microbiol 8:997.

4. Baker-Austin C, Trinanes JA, Taylor NGH, Hartnell R, Siitonen A, Martinez-Urtaza J. 2013. Emerging Vibrio risk at high latitudes in response to ocean warming. 1. Nature Clim Change 3:73–77.

5. Vezzulli L, Colwell RR, Pruzzo C. 2013. Ocean Warming and Spread of Pathogenic Vibrios in the Aquatic Environment. Microb Ecol 65:817–825.

6. Vezzulli L, Colwell RR, Pruzzo C. 2003. Emerging issues in water and infectious disease. World Health Organization, Geneva.

7. Archer EJ, Baker-Austin C, Osborn TJ, Jones NR, Marinez-Urtaza J, Trinanes J, Oliver JD, González FJC, Lake IR. 2023. Climate warming and increasing Vibrio vulnificus infections in North America. 1. Sci Rep 13:3893.

8. Iwamoto M, Ayers T, Mahon BE, Swerdlow DL. 2010. Epidemiology of Seafood-Associated Infections in the United States. Clin Microbiol Rev 23:399.

9. Newton A, Kendall M, Vugia DJ, Henao OL, Mahon BE. 2012. Increasing rates of vibriosis in the United States, 1996–.2010: review of surveillance data from 2 systems. Clin Infect Dis 54:391–395.

10. Brumfield KD, Chen AJ, Gangwar M, Usmani M, Hasan NA, Jutla AS, Huq A, Colwell RR. 2023. Environmental Factors Influencing Occurrence of Vibrio parahaemolyticus and Vibrio vulnificus. Appl Environ Microbiol 89:e00307–23.

11. Slayton RB, Newton AE, DePaola A, Jones JL, Mahon BE. 2014. Clam-associated vibriosis. Infect 142:1083–1088.

12. Wallace BJ, Guzewich JJ, Cambridge M, Altekruse S, Morse DL. 1999. Seafood-associated disease outbreaks in New York, 1980-1994. Am J Prev Med 17:48–54.

13. Wechsler E, D’Aleo C, Hill VA, Hopper J, Myers-Wiley D, O’Keeffe E, Jacobs J, Guido F, Huang A, Dodt SN, Rowan B, Sherman M, Greenberg A, Schneider D, Noone B, Fanella L, Williamson BR, Dinda E, Mayer M, Backer M, Agasan A, Kornstein L, Stavinsky F, Neal B, Edwards D, Haroon M, Hureley D, Cobert L, Miller J, Mojica B, Carloni E, Devine B, Cambridge M, Root T, Schoonmaker D, Shayegani M, Hastback E, Wallace B, Kondracki S, Smith P, Matiuck S, Pilot K, Acharya M, Wolf G, Manley W, Genese C, Brooks J, Hadler J. 1999. Outbreak of Vibrio parahaemolyticus infection associated with eating raw oysters and clams harvested from Long Island Sound. Mortal Wkly Rep 48:48–51.

14. Vezzulli L, Pruzzo C, Huq A, Colwell RR. 2010. Environmental reservoirs of Vibrio cholerae and their role in cholera: Environmental reservoirs of V. cholerae. Environmental Microbiology Reports 2:27– 33.

15. Pruzzo C, Vezzulli L, Colwell RR. 2008. Global impact of Vibrio cholerae interactions with chitin. Environ Microbiol 10:1400–1410.

16. Colwell RR. 1996. Global Climate and Infectious Disease: The Cholera Paradigm. Science 274:2025– 2031.

17. Cook DW, Bowers JC, DePaola A. 2002. Density of total and pathogenic (tdh+) Vibrio parahaemolyticus in Atlantic and Gulf coast molluscan shellfish at harvest. J Food Prot 65:1873– 1880.

18. DePaola A, Nordstrom JL, Bowers JC, Wells JG, Cook DW. 2003. Seasonal abundance of total and pathogenic Vibrio parahaemolyticus in Alabama oysters. Appl Environ Microbiol 69:1521–1526.

19. Oliver JD. 1989. Vibrio vulnificus, p. 569–600. In Doyle, MP (ed.), Foodborne bacterial pathogens. Marcel Dekker, Inc, New York.

20. Kaneko T, Colwell RR. 1973. Ecology of <VIBRIO parahaemolyticus> in Chesapeake Bay. J Bacteriol 113:24.

21. Wright AC. 1993. Rapid identification of Vibrio vulnificus on nonselective media with an alkaline phosphatase-labeled oligonucleotide probe. Appl Environ Microbiol 59:541–546.

22. Banakar V, Magny G C, J J, R M, A H, RJ W, RR C. 2011. Temporal and spatial variability in the distribution of Vibrio vulnificus in the Chesapeake Bay: a hindcast study. Ecohealth 8:456–467.

23. Socioeconomic Value of the Chesapeake Bay Watershed in Delaware. https://udspace.udel.edu/items/d7334b54-a6c5-4693-a572-8004206bc25f. Retrieved 14 September 2023.

24. Corrozi Narvaez M, Kauffman G. 2012. Economic Benefits and Jobs Provided by Delaware Watersheds.

25. Blackwell KD, Oliver JD. 2008. The ecology of Vibrio vulnificus, Vibrio cholerae, and Vibrio parahaemolyticus in North Carolina Estuaries. J Microbiol 46:146–153.

26. Grimes DJ, Johnson CN, Dillon KS, Flowers AR, Noriea NF, Berutti T. 2009. What Genomic Sequence Information Has Revealed About Vibrio Ecology in the Ocean—A Review. Microb Ecol 58:447–460.

27. Zimmerman AM, DePaola A, Bowers JC, Krantz JA, Nordstrom JL, Johnson CN, Grimes DJ. 2007. Variability of Total and Pathogenic Vibrio parahaemolyticus Densities in Northern Gulf of Mexico Water and Oysters. Applied and Environmental Microbiology 73:7589–7596.

28. Igbinosa EO, Obi CL, Okoh AI. 2011. Seasonal abundance and distribution of Vibrio species in the treated effluent of wastewater treatment facilities in suburban and urban communities of Eastern Cape Province, South Africa. J Microbiol 49:224–232.

29. Parveen S, Hettiarachchi KA, Bowers JC, Jones JL, Tamplin ML, McKay R, Beatty W, Brohawn K, DaSilva LV, DePaola A. 2008. Seasonal distribution of total and pathogenic Vibrio parahaemolyticus in Chesapeake Bay oysters and waters. International Journal of Food Microbiology 128:354–361.

30. Ramirez GD, Buck GW, Smith AK, Gordon KV, Mott JB. 2009. Incidence of *Vibrio vulnificus* in estuarine waters of the south Texas Coastal Bend region. Journal of Applied Microbiology 107:2047–2053.

31. Caburlotto G, Haley BJ, Lleò MM, Huq A, Colwell RR. 2010. Serodiversity and ecological distribution of Vibrio parahaemolyticus in the Venetian Lagoon, Northeast Italy. Environmental Microbiology Reports 2:151–157.

32. Deter J, Lozach S, Derrien A, Véron A, Chollet J, Hervio-Heath D. 2010. Chlorophyll a might structure a community of potentially pathogenic culturable Vibrionaceae. Insights from a one-year study of water and mussels surveyed on the French Atlantic coast. Environmental Microbiology Reports 2:185–191.

33. Johnson CN, Flowers AR, Noriea NF III, Zimmerman AM, Bowers JC, DePaola A, Grimes DJ. 2010. Relationships between environmental factors and pathogenic vibrios in the Northern Gulf of Mexico. Appl Environ Microbiol76:7076–7084Johnson et al.

34. Asplund ME, Rehnstam-Holm A-S, Atnur V, Raghunath P, Saravanan V, Härnström K, Collin B, Karunasagar I, Godhe A. 2011. Water column dynamics of Vibrio in relation to phytoplankton community composition and environmental conditions in a tropical coastal area. Environmental Microbiology 13:2738–2751.

35. Martinez-Urtaza J, Blanco-Abad V, Rodriguez-Castro A, Ansede-Bermejo J, Miranda A, Rodriguez-Alvarez MX. 2012. Ecological determinants of the occurrence and dynamics of Vibrio parahaemolyticus in offshore areas. 5. ISME J 6:994–1006.

36. Occurrence of Vibrio vulnificus Biotypes in Danish Marine Environments | Applied and Environmental Microbiology. https://journals.asm.org/doi/full/10.1128/AEM.64.1.7-13.1998. Retrieved 11 October 2022.

37. Randa MA, Polz MF, Lim E. 2004. Effects of Temperature and Salinity on Vibrio vulnificus Population Dynamics as Assessed by Quantitative PCR. Applied and Environmental Microbiology 70:5469– 5476.

38. Wright AC, Hill RT, Johnson JA, Roghman MC, Colwell RR, JG M Jr. 1996. Distribution of Vibrio vulnificus in the Chesapeake Bay. Appl Environ Microbiol 62:717–724.

39. Kaneko T, Colwell RR. 1975. Incidence of Vibrio parahaemolyticus in Chesapeake Bay. Appl Microbiol 30:251–257.

40. Kaspar CW, Tamplin ML. 1993. Effects of temperature and salinity on the survival of Vibrio vulnificus in seawater and shellfish. Appl Environ Microbiol 59:2425–2429.

41. Oliver JD. 2015. The Biology of *Vibrio vulnificus*. Microbiol Spectr 3:3.3.01.

42. Kelly MT. 1982. Effect of temperature and salinity on Vibrio (Beneckea) vulnificus occurrence in a Gulf Coast environment. Appl Environ Microbiol 44:820–824.

43. Lin M, Payne DA, Schwarz JR. 2003. Intraspecific Diversity of Vibrio vulnificus in Galveston Bay Water and Oysters as Determined by Randomly Amplified Polymorphic DNA PCR. Applied and Environmental Microbiology 69:3170–3175.

44. Predicting the distribution of Vibrio vulnificus in Chesapeake Bay. https://repository.library.noaa.gov/view/noaa/2531. Retrieved 25 August 2023.

45. Audemard C, Ben-Horin T, Kator HI, Reece KS. 2022. Vibrio vulnificus and Vibrio parahaemolyticus in Oysters under Low Tidal Range Conditions: Is Seawater Analysis Useful for Risk Assessment? 24. Foods 11:4065.

46. Lipp EK, Rodriguez-Palacios C, Rose JB. 2001. Occurrence and distribution of the human pathogen Vibrio vulnificus in a subtropical Gulf of Mexico estuary, p. 165–173. In Porter, JW (ed.), The Ecology and Etiology of Newly Emerging Marine Diseases. Springer Netherlands, Dordrecht.

47. Wetz JJ, AD B, JS F, ZF W, RT N. 2014. Quantification of Vibrio vulnificus in an Estuarine Environment: a Multi-Year Analysis Using QPCR. Estuaries and Coasts 37:421–435.

48. Pfeffer CS, Hite MF, Oliver JD. 2003. Ecology of Vibrio vulnificus in Estuarine Waters of Eastern North Carolina. Appl Environ Microbiol 69:3526–3531.

49. Böer SI, Heinemeyer E-A, Luden K, Erler R, Gerdts G, Janssen F, Brennholt N. 2013. Temporal and Spatial Distribution Patterns of Potentially Pathogenic Vibrio spp. at Recreational Beaches of the German North Sea. Microb Ecol 65:1052–1067.

50. Esteves K, Hervio-Heath D, Mosser T, Rodier C, Tournoud M-G, Jumas-Bilak E, Colwell RR, Monfort P. 2015. Rapid Proliferation of Vibrio parahaemolyticus, Vibrio vulnificus, and Vibrio cholerae during Freshwater Flash Floods in French Mediterranean Coastal Lagoons. Applied and Environmental Microbiology 81:7600–7609.

51. Høi L, Larsen JL, Dalsgaard I, Dalsgaard A. 1998. Occurrence of Vibrio vulnificus Biotypes in Danish Marine Environments. Applied and Environmental Microbiology 64:7–13.

52. Rodriguez-Castro A, Ansede-Bermejo J, Blanco-Abad V, Varela-Pet J, Garcia-Martin O, Martinez-Urtaza J. 2010. Prevalence and genetic diversity of pathogenic populations of Vibrio parahaemolyticus in coastal waters of Galicia, Spain. Environmental Microbiology Reports 2:58–66.

53. Davis BJK, Jacobs JM, Davis MF, Schwab KJ, DePaola A, Curriero FC. 2017. Environmental Determinants of Vibrio parahaemolyticus in the Chesapeake Bay. Appl Environ Microbiol 83:e01147–17.

54. Paranjpye RN, WB N, M L, ED H, BJ G, Q L, BD B, VL T, MS S, PA S. 2015. Environmental influences on the seasonal distribution of Vibrio parahaemolyticus in the Pacific Northwest of the USA. FEMS Microbiology Ecology 91:1–11.

55. Julie D, Solen L, Antoine V, Jaufrey C, Annick D, Dominique H-H. 2010. Ecology of pathogenic and non-pathogenic Vibrio parahaemolyticus on the French Atlantic coast. Effects of temperature, salinity, turbidity and chlorophyll a. Environmental Microbiology 12:929–937.

56. Sarkar BL, Nair GB, Sircar BK, Pal SC. 1983. Incidence and level of Vibrio parahaemolyticus associated with freshwater plankton. Appl Environ Microbiol 46:288–290.

57. Sarkar BL, Nair GB, Banerjee AK, Pal SC. 1985. Seasonal distribution of Vibrio parahaemolyticus in freshwater environs and in association with freshwater fishes in Calcutta. Appl Environ Microbiol 49:132–136.

58. Patra AK, Acharya BC, Mohapatra A. 2009. Occurrence and distribution of bacterial indicators and pathogens in coastal waters of Orissa. IJMS Vol38(4) [December 2009].

59. Rudas T. 1998. Odds ratios in the analysis of contingency tables. Sage.

60. Racault M-F, Abdulaziz A, George G, Menon N, C J, Punathil M, McConville K, Loveday B, Platt T, Sathyendranath S, Vijayan V. 2019. Environmental Reservoirs of Vibrio cholerae: Challenges and Opportunities for Ocean-Color Remote Sensing. 23. Remote Sensing 11:2763.

61. Vezzulli L. 2022. Global expansion of Vibrio spp. in hot water. Environmental Microbiology Reports 15:77–79.

62. Parveen S, Jacobs J, Ozbay G, Chintapenta LK, Almuhaideb E, Meredith J, Ossai S, Abbott A, Grant A, Brohawn K. 2020. Seasonal and geographical differences in total and pathogenic Vibrio parahaemolyticus and Vibrio vulnificus levels in seawater and oysters from the Delaware and Chesapeake Bays determined using several methods. Applied and environmental microbiology 86:e01581–20.

63. Wise BM, NB G. 1996. The process chemometrics approach to process monitoring and fault detection. Journal of Process Control 6:329–348.

64. McNutt L-A, Wu C, Xue X, Hafner JP. 2003. Estimating the relative risk in cohort studies and clinical trials of common outcomes. American journal of epidemiology 157:940–943.

65. Schmidt CO, Kohlmann T. 2008. When to use the odds ratio or the relative risk? International journal of public health 53:165.

66. Davis BJ, Corrigan AE, Sun Z, Atherly E, DePaola A, Curriero FC. 2021. A case-control analysis of traceback investigations for Vibrio parahaemolyticus infections (vibriosis) and pre-harvest environmental conditions in Washington State, 2013–2018. Science of the Total Environment 752:141650.

67. Onohuean H, Okoh AI, Nwodo UU. 2021. Epidemiologic potentials and correlational analysis of Vibrio species and virulence toxins from water sources in greater Bushenyi districts, Uganda. Scientific reports 11:1–16.

